# CRISPR-Cas immunity repressed by a biofilm-activating pathway in *Pseudomonas aeruginosa*

**DOI:** 10.1101/673095

**Authors:** Adair L. Borges, Bardo Castro, Sutharsan Govindarajan, Tina Solvik, Veronica Escalante, Joseph Bondy-Denomy

## Abstract

CRISPR-Cas systems are adaptive immune systems that protect bacteria from bacteriophage (phage) infection. To provide immunity, RNA-guided protein surveillance complexes recognize foreign nucleic acids, triggering their destruction by Cas nucleases. While the essential requirements for immune activity are well understood, the physiological cues that regulate CRISPR-Cas expression are not. Here, a forward genetic screen identifies a two-component system (KinB/AlgB), previously characterized in regulating *Pseudomonas aeruginosa* virulence and biofilm establishment, as a regulator of the biogenesis and activity of the Type I-F CRISPR-Cas system. Downstream of the KinB/AlgB system, activators of biofilm production AlgU (a σ^E^ orthologue) and AlgR, act as repressors of CRISPR-Cas activity during planktonic and surface-associated growth. AmrZ, another biofilm activator, functions as a surface-specific repressor of CRISPR-Cas immunity. *Pseudomonas* phages and plasmids have taken advantage of this regulatory scheme, and carry hijacked homologs of AmrZ, which are functional CRISPR-Cas repressors. This suggests that while CRISPR-Cas regulation may be important to limit self-toxicity, endogenous repressive pathways represent a vulnerability for parasite manipulation.

---

Clustered regularly interspaced short palindromic repeats (CRISPR) and CRISPR-associated (cas) genes are RNA-guided nucleases found in nearly half of all bacteria^1^. CRISPR-Cas systems are mechanistically diverse, with six distinct types (I-VI) identified, based on signature genes, mechanisms of action, and the type of nucleic acid target^1^. Our strong understanding of the basic components to enable sequence specific DNA and RNA cleavage have enabled functional transplantation into heterologous bacterial^2,3^ and eukaryotic^4,5^ hosts. Given that many Cas proteins encode nucleases^6^ the fine-tuned regulation of these systems to avoid toxicity is likely a key factor to enable safe retention of a CRISPR-Cas immune system^7^. Multiple signals have been shown to activate CRISPR-Cas function in diverse organisms, such as quorum sensing^8,9^, temperature^10^, membrane stress^11,12^, altered host metabolite levels^13,14^, and phage infection^15-18^. However, relatively little is known regarding the factors and/or signals that serve to temper CRISPR-Cas activity and mitigate the risk of acquiring and expressing a nucleolytic immune system.

Type I CRISPR-Cas systems are comprised of a multi-subunit RNA-guided surveillance complex, a trans-acting nuclease (Cas3)^19-21^, and proteins dedicated to spacer acquisition, Cas1 and Cas2^22,23^. *Pseudomonas aeruginosa* has become a powerful model organism for studying Type I CRISPR-Cas mechanisms^24-29^, functions^3,23,30,31^, evolution^32-34^, and interactions with phages utilizing anti-CRISPR proteins^35-38^. The *P. aeruginosa* strain PA14 possesses an active Type I-F CRISPR-Cas immune system that naturally targets many phages. The surveillance complex is composed of four Cas proteins (Csy1, Csy2, Csy3, and Csy4), which are encoded together in a single operon^26^. The surveillance complex is guided by CRISPR RNAs (crRNAs) originating from one of two chromosomally encoded CRISPR arrays^26^. Finally, a separate operon is responsible for the production of Cas1 and a Cas2-3 fusion protein, the latter of which mediates nucleolytic destruction of target DNA.

To discover regulators controlling the expression of the CRISPR-Cas surveillance complex in *P. aeruginosa*, we utilized a *P. aeruginosa* strain PA14 engineered to express *lacZ* in place of the *csy3* gene (*csy3::lacZ*)^39^. The *csy3*::*lacZ* reporter strain was subjected to transposon mutagenesis and colonies isolated and screened on X-gal plates. ∼40,000 colonies were visually examined for increased or decreased levels of β-galactosidase. Multiple independent insertions were identified that abolished LacZ levels within *lacZ* and upstream genes (*csy1* and *csy2*), indicating successful mutagenesis and screening. Thirty distinct mutants with transposon insertions outside of this region, with altered β-galactosidase levels, were isolated and mapped. These mutants are listed in Supplementary Table 1 with β-galactosidase levels quantified for mutants that grew well in culture. Four independent insertions were identified in a single gene, *kinB*, which resulted in apparent mucoidy plating morphology and decreased β-galactosidase production (Supplementary Fig. 1a). Quantification of β-galactosidase levels in liquid growth experiments revealed ∼50% less *csy3::lacZ* expression compared to wild-type (WT) PA14 for each mutant (Supplementary Table 1, Supplementary Fig. 1b). We focused on *kinB* (a sensor kinase/phosphatase) because it was the only gene with >1 independent transposon insertion and displayed the largest β-galactosidase activity change.

To determine the consequences of *kinB* disruption and decreased *csy* expression on CRISPR-Cas function, we measured the ability of *kinB::Tn* insertions to limit the replication of phage, when introduced into a wild-type strain background. Phages used to assay activity are: DMS3_*acrIE3*_ which is an untargeted control phage, DMS3m_*acrIE3*_ ^40^, which is fully targeted by the PA14 Type I-F CRISPR-Cas system, and phage DMS3m_*acrIF4*_, which is partially targeted, by virtue of encoding a “weak” anti-CRISPR, *acrIF4*, that binds to the surveillance complex to inhibit CRISPR-Cas function^35,37,41^. The *kinB::Tn* strains remained resistant to DMS3m_*acrIE3*_ infection, but DMS3m_*acrIF4*_ formed 10-fold more plaques relative to WT, demonstrating attenuated CRISPR-Cas function (Figure 1a, Supplementary Fig. 1c). This defect was complemented by expression of *kinB in trans* (Supplementary Fig. 1c). Growth of control phage DMS3_*acrIE3*_ was not impacted in the absence of KinB (Figure 1a). Furthermore, two other targeted phages, JBD26 (naturally possessing *acrIF4*) and JBD25 (a partially targeted phage with no Acr) also showed increased replication in the *kinB:Tn* strain (Supplementary Fig. 1d) relative to WT PA14. Replication of a phage with a weak anti-CRISPR or one that is targeted by a less active spacer is therefore a sensitive barometer for perturbations in CRISPR-Cas levels. Together, these data confirm that in the absence of *kinB, csy* gene expression and phage targeting are decreased.

**Figure 1:**
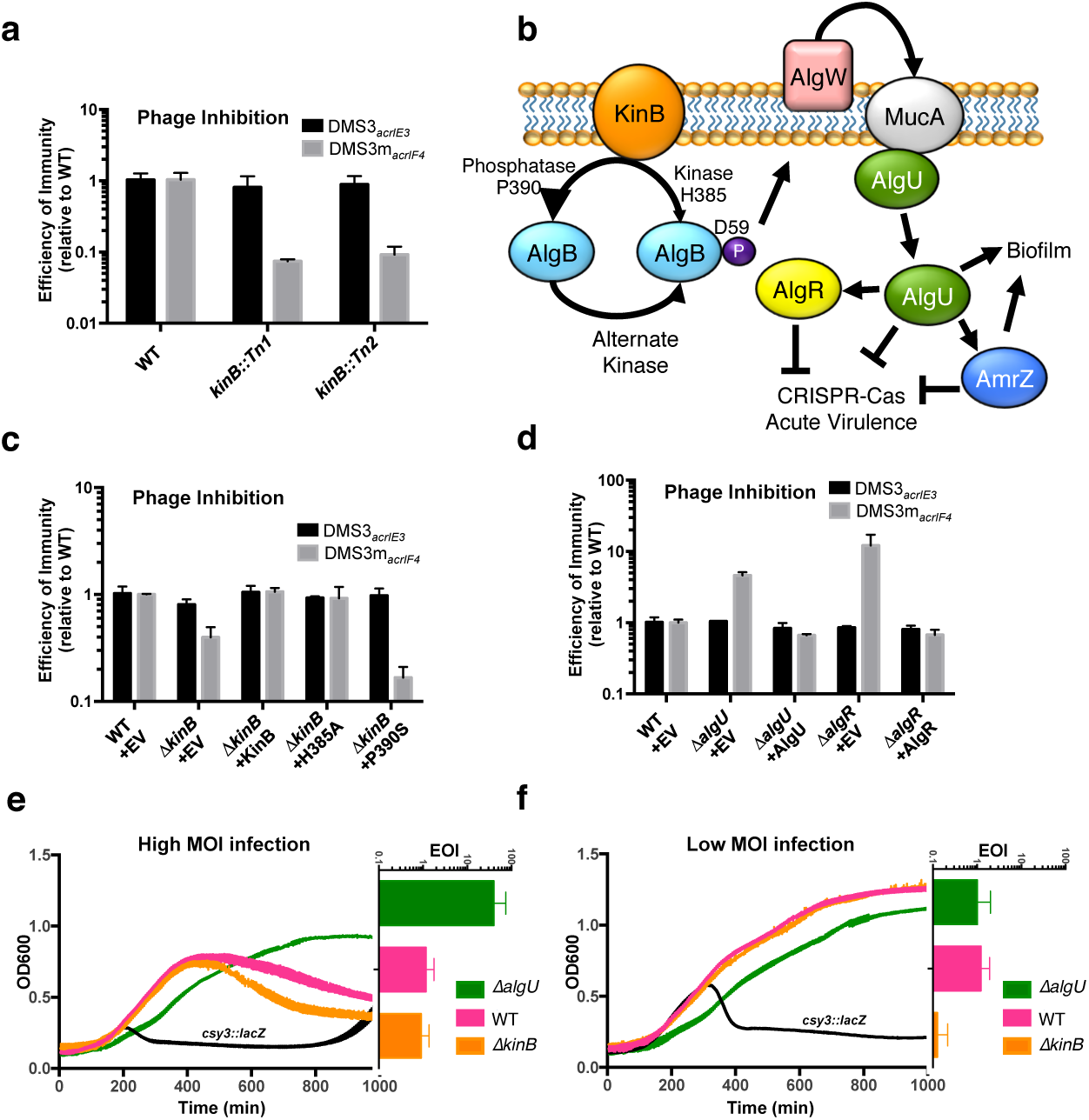
A forward genetic screen identifies a role for a biofilm activating pathway in repressing CRISPR-Cas immunity. **a**. Efficiency of immunity (EOI) against isogenic phages DMS3_*acrIE3*_ (non-targeted) and DMS3m_*acrIF4*_ (CRISPR-targeted). Plaque forming units (PFUs) were quantified on two independent *kinB* transposon mutants (*kinB::Tn1* and *kinB::Tn2*), then represented as a ratio relative to the number of PFUs measured on WT PA14. **b.** A cartoon summarizing the KinB/AlgB two component system and downstream effects, based on prior work, with CRISPR-Cas regulation added. **c**,**d** EOI measurements for indicated *ΔkinB, ΔalgR*, and *ΔalgU* strains with complementation. **e**,**f**. Growth curves of PA14 WT, *ΔalgU*, or *ΔkinB* strains infected with either a high (e,10^6^ PFU) or low (f, 10^4^ PFU) MOI of virulent DMS3m_*acrIF4*_. Phage replication was quantified as PFUs after 24 hours, and the efficiency of immunity expressed as a ratio of the number of PFUs harvested from the mutant strain compared to the number of PFUs obtained from WT. All data (OD600 and EOI measurements) are represented as the mean of 3 biological replicates +/-SD.

KinB is a sensor kinase/phosphatase in a two-component system with response regulator AlgB. The KinB/AlgB system has a large regulon within *P. aeruginosa*, but is well-known for its role in biofilm regulation^42^. In the absence of KinB function, the activity of the transcription factor AlgU (a σ^E^ orthologue and an activator of biofilm formation) is enhanced^43,44^. This activation of AlgU manifests solely due to the lack of KinB phosphatase activity and the resulting accumulation of the phosphorylated form of the response regulator AlgB (P-AlgB). The phosphorylation of AlgB has been attributed to unknown kinases^45,46^ (Fig. 1b). The resulting accumulation of P-AlgB activates the periplasmic protease AlgW (a DegS homolog), which degrades MucA, liberating sigma factor AlgU^47,48,49^ (Fig. 1b). AlgU positively regulates many factors involved in biofilm production, including AlgR, AlgD, AlgB, and AmrZ^50-52^.

To determine if this KinB-controlled biofilm activation pathway also regulates CRISPR-Cas immunity, we tested the ability of phosphatase and kinase dead KinB mutants to complement an in-frame *ΔkinB* deletion, using the replication of the partially-targeted DMS3m_*acrIF4*_ phage as a metric for CRISPR-Cas activity. WT *kinB* or kinase inactive H385A *kinB* complemented the CRISPR defect (Fig. 1c), indicating KinB kinase activity is dispensable for CRISPR-Cas activation. However, a P390S *kinB* mutant incapable of dephosphorylating the response regulator AlgB did not complement, and in fact decreased CRISPR-Cas activity further (Fig. 1c). This suggests that CRISPR-Cas expression is inhibited by the accumulation of P-AlgB. A Δ*kinBΔalgB* double mutant restored CRISPR-Cas targeting to WT levels (Supplementary Fig. 2a), confirming the role of this signaling pathway. A strain lacking *algB* (*ΔalgB*) or possessing a D59N mutant that cannot be phosphorylated had elevated CRISPR-Cas activity, supporting the repressive role of P-AlgB (Supplementary Fig. 2b). Together, these data suggest that KinB-mediated dephosphorylation of AlgB lifts CRISPR-Cas inhibition, and conversely accumulation of high levels of P-AlgB (achieved in Δ*kinB* or *kinB P390S*) leads to CRISPR-Cas repression.

To assess CRISPR-Cas regulation downstream of P-AlgB, anti-phage immunity was assayed in *ΔalgU* and *ΔalgR* backgrounds, revealing enhanced targeting of DMS3m_*acrIF4*_ but not control phage DMS3_*acrIE3*_ in each knockout (Fig. 1d). Complementation with *algR* and *algU* restored CRISPR-Cas levels in their respective knockout backgrounds (Fig. 1d). This demonstrates that these proteins repress CRISPR-Cas immunity. Double knockouts of each gene combined with *ΔkinB* each resembled the respective *ΔalgU* and *ΔalgR* single knockouts, consistent with these factors acting downstream of KinB (Supplementary Fig. 2a). The observed plaquing defects seen with the DMS3m_*acrIF4*_ phage are CRISPR-dependent, as double knockouts (*kinB, algB, algU, algR* mutants combined with *csy3::lacZ*) revealed plaquing equivalent to *csy3::lacZ* alone (Supplementary Fig. 2c).

Phages that use anti-CRISPRs to inhibit CRISPR-Cas immunity must cooperatively infect the same cell in order to produce a sufficient dose of anti-CRISPR to permit phage replication^41,53^. The multiplicity of infection (MOI) required to overwhelm immunity is dependent on the affinity of the anti-CRISPR for the CRISPR-Cas machinery and the intracellular concentration of CRISPR-Cas components. To understand how this regulatory pathway controls phage cooperation requirements on a population level by potentially shifting the intracellular concentration of CRISPR-Cas complexes, liquid cultures of *ΔalgU* and *ΔkinB* mutants were infected with a virulent derivative of phage DMS3m_*acrIF4*_ at both high (where the WT cells are sensitive) and low (where the WT cells are resistant) MOI (Figure 1e). The *ΔalgU* mutant was able to limit phage replication relative to WT and continue growing when infected with concentration of phages sufficiently high to overwhelm WT immunity (10^6^ PFU), indicating increased levels of CRISPR-Cas machinery (Fig 1e). Conversely, when infected with a low MOI of phages (10^4^ PFU), all bacterial genotypes grew to high cell density, however the *ΔkinB* mutant permitted increased phage replication relative to WT, demonstrating a decreased level of immune system components (Fig. 1f). Importantly, these changes in phage replication requirements were not seen in the absence of CRISPR-Cas immunity, as phage replication in double knockouts *algU csy3::lacZ and kinB csy3::lacZ* resembles *csy3::lacZ (*Supplementary Fig. 3a-d, f). In contrast, *ΔalgR* demonstrated CRISPR-independent phage resistance during liquid culture infection, and phage cooperation requirements could not be reliably measured for this mutant (Supplementary Fig. 3e). Taken together, these data demonstrate that a well-characterized biofilm activation pathway controlled by alternate sigma factor, AlgU, represses CRISPR-Cas immunity in *P. aeruginosa.*

Our screen focused on finding regulators of the *csy* operon, however the recruited Cas3 nuclease (encoded in a separate, nearby operon) is also an important player in Type I CRISPR-Cas immunity. To measure the relative impact of the KinB/AlgB pathway in controlling Cas3 and Csy complex levels, we fused an N-terminal sfCherry tag to the endogenous *csy1* or *cas3* gene in the mutant backgrounds. We observed decreased levels of both Cas3-sfCherry and Csy1-sfCherry in *ΔkinB* and increased levels in *ΔalgR* and *ΔalgU* relative to WT, demonstrating that this pathway controls the levels of both Cas3 and the Csy complex in the bacterial cell (Fig. 2a).

**Figure 2.**
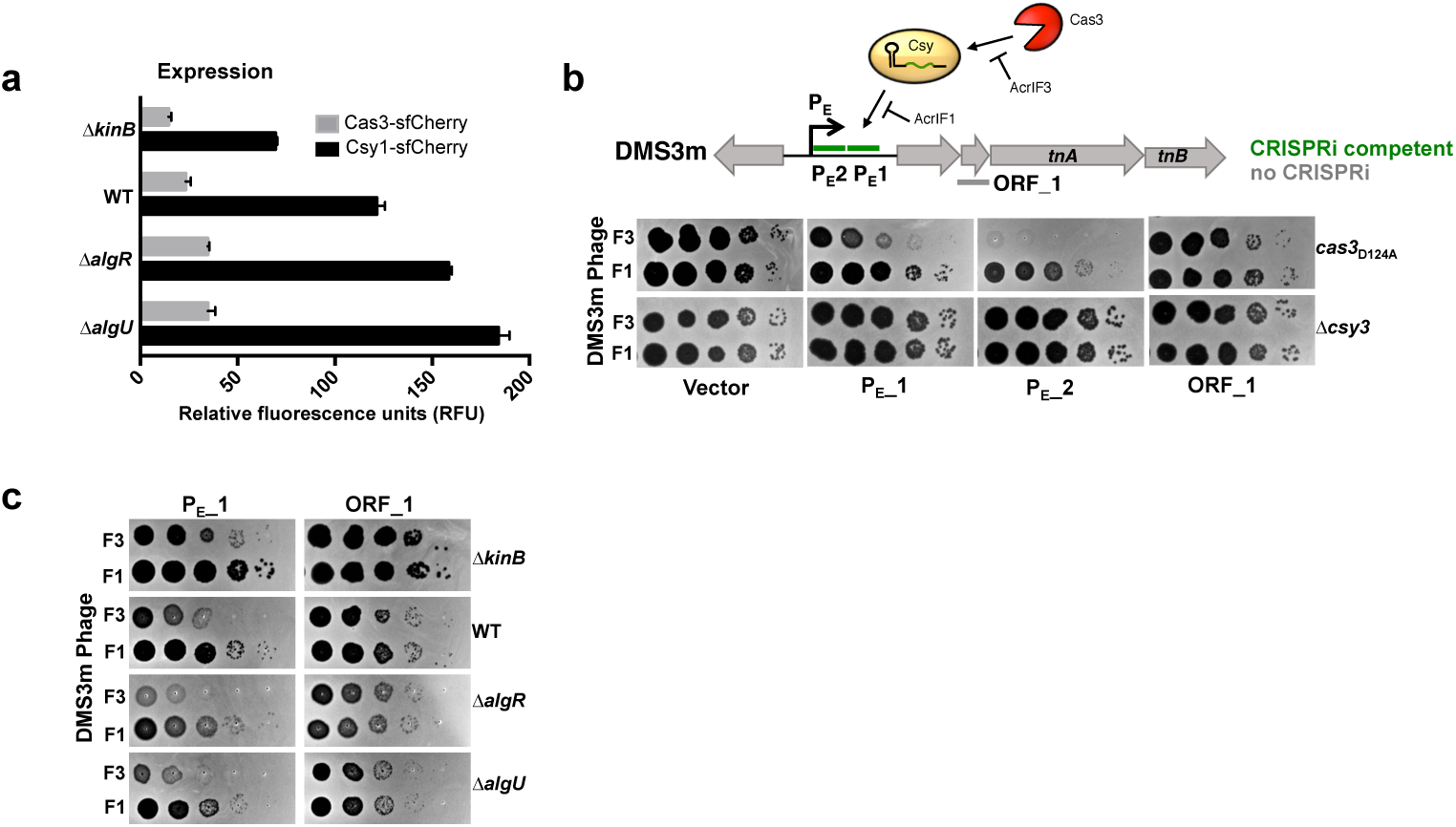
The kinB/algB pathway modulates Csy complex levels and activity. **a.**Measurement of the fluorescence levels of Cas3-sfCherry (grey) or Csy1-sfCherry (black) reporter strains after 10 hours of growth in liquid culture. Fluorescence measurements are represented as the mean of 3 biological replicates +/-SD. **b**. Spot titration of DMS3m_*acrIF3*_ (F3) and DMS3m_*acrIF1*_ (F1) on *cas3*_D124A_ (dead Cas3) or Δcsy3 (active Cas3, no Csy complex). Phages are targeted by natural spacer CR2_1, as well as crRNAs designed to target DMS3m genome in positions designated on ORF map. **c**. Spot titration of DMS3m_*acrIF3*_ and DMS3m_*acrIF1*_ phages on deletion mutants expressing the indicated crRNA.

As Cas3-sfCherry was expressed at low levels relative to Csy1-sfCherry, and is also known to be subject to post-translational control by Cas1^28^, we sought to dissect the relative contribution of nuclease versus surveillance complex disregulation in driving the immune phenotypes of the KinB/AlgB pathway mutants. To specifically measure the anti-phage activity of the Csy complex, we developed a Cas3-indepedent bioassay to read out the activity of the surveillance complex in the cell. Through the rational design of crRNAs to target an early phage promoter (P_E__1, P_E__2), we observed inhibition of phage replication in a *P. aeruginosa* strain with a nuclease dead Cas3 (Cas3_D124A_, Fig. 2a). This CRISPR-based transcriptional interference (CRISPRi) effect was also seen when infecting with a phage that expressed the inhibitor of Cas3 recruitment, AcrIF3, but not an inhibitor that blocks Csy complex-phage DNA binding^37^, AcrIF1 (Fig. 2b). CRISPRi was mediated by crRNAs targeting phage promoters, including one (P_E__2) that completely limited phage replication in the absence of Cas3 activity, while an ORF-targeting crRNA (ORF_1) was ineffective (Fig. 2b). We selected P_E__1 as a moderately-functional CRISPRi spacer and expressed it in various mutant backgrounds generated in this study. We observed decreased CRISPRi activity against phage DMS3m_*acrIF3*_ in the ΔkinB background, but increased CRISPRi-mediated phage repression in *ΔalgR* and *ΔalgU*, (Fig. 2c), demonstrating that modulation of *csy* gene expression by this pathway is sufficient to impact phage targeting, in a Cas3-independent manner. We conclude that while the KinB/AlgB pathway regulates both Cas3 and Csy complex levels, repression of surveillance complex biogenesis alone represents a powerful mechanism by which the KinB/AlgB pathway controls CRISPR-Cas immunity.

AlgU and AlgR are proteins that regulate hundreds of genes in *P. aeruginosa*^54-56^, ultimately repressing acute virulence factors (e.g. pyocyanin) and activating chronic virulence factors (e.g. biofilm production). To identify downstream CRISPR-Cas regulators, we considered the large AlgU regulon^56^, but focused on a factor involved in biofilm production, whose expression is AlgU-dependent, *amrZ*^51,57^. AmrZ is a ribbon-helix-helix type transcription factor, and acts as both a repressor and activator of transcription in *Pseudomonas*^58^. We generated a knockout of *amrZ* and observed a CRISPR-dependent decrease in DMS3m_*acrIF4*_ plaque formation (Fig. 3a, Supplementary Fig. 2c). This was complemented when *amrZ* was expressed *in trans* (Fig. 3a), indicating that AmrZ is also a repressor of CRISPR-Cas immunity. However, when we measured phage cooperation requirements in the *ΔamrZ* strain, phage replication and immune protection did not differ substantially from WT (Fig. 3b,c). The most obvious difference between the phage plaque assay and the assay to measure phage cooperation is that a plaque assay is performed on solid plates whereas phage cooperation requirements were measured in liquid culture. To test if AmrZ was a surface-specific repressor of CRIPSR-Cas immunity, we measured the levels of Csy complex during surface association and planktonic growth in WT and *ΔamrZ* cells using an endogenous *csy1-sfCherry* reporter. In WT cells, the levels of Csy complex were attenuated during surface-association relative to planktonic growth, but in the absence of AmrZ, Csy complex levels during surface association increased to levels comparable to those in planktonic growth (Fig. 3d). Deletion of *amrZ* did not impact Csy complex levels during planktonic growth (Fig. 3e). To artificially increase the levels of AmrZ during planktonic growth, we ectopically expressed AmrZ from a high copy plasmid. In this scenario, high levels of AmrZ were sufficient to completely inhibit Csy complex expression, suggesting that low AmrZ activity in planktonic growth underlies its surface-specific control of CRISPR-Cas (Fig. 3f). In contrast to AmrZ, overexpression of the AlgU moderately impacted Csy complex levels and AlgR was unable to repress *csy1-sfCherry* when overexpressed (Fig. 3f). This is likely because AlgU and AlgR are master regulators and many redundant mechanisms are in place to control their activity. Taken together, these data demonstrate that AmrZ is a surface-specific repressor of the *csy* operon.

**Figure 3:**
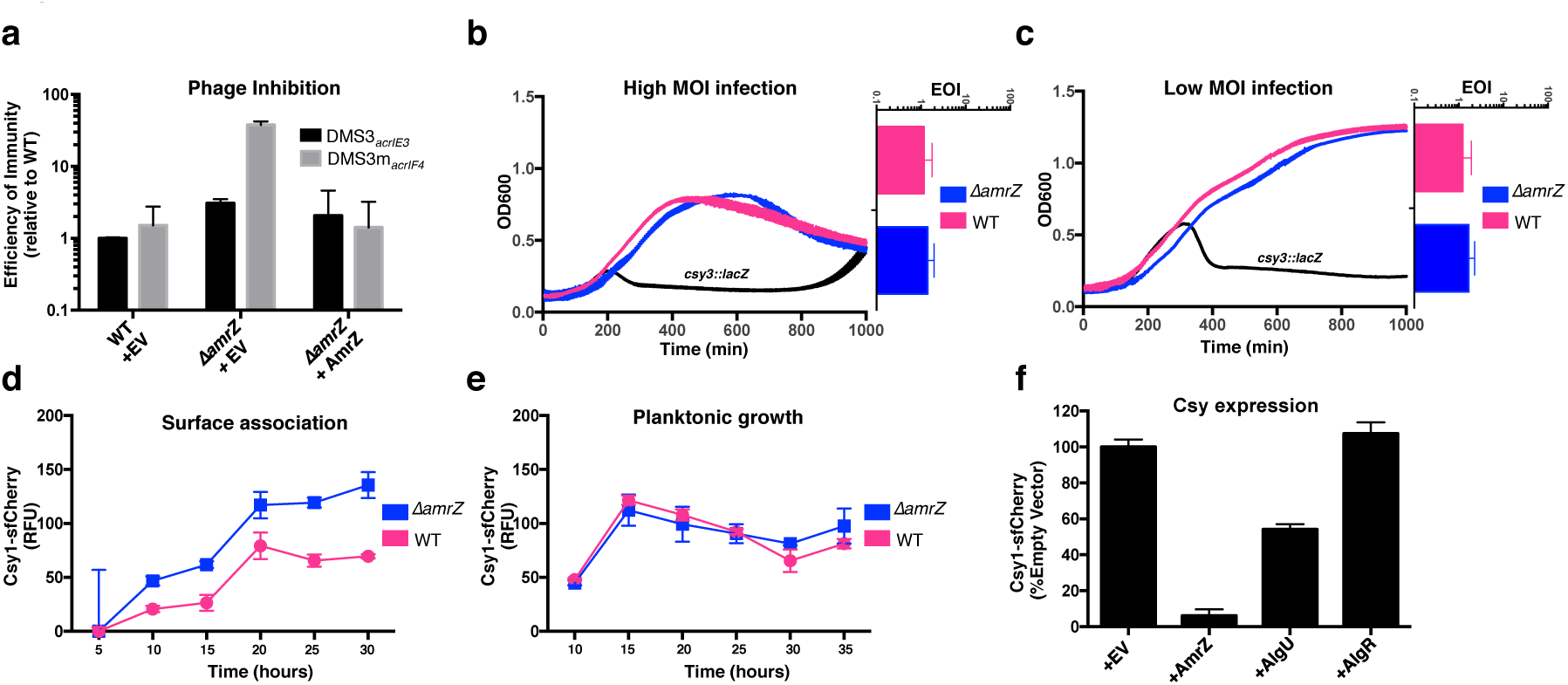
AmrZ is a surface-specific repressor of CRISPR-Cas immunity. **a**. Efficiency of immunity (EOI) against isogenic phages DMS3_*acrIE3*_ (non-targeted) and DMS3m_*acrIF4*_ (CRISPR-targeted). Plaque forming units (PFUs) were quantified on ΔamrZ or the complemented strain, then represented as a ratio of the number of PFUs measured on WT PA14. EOI measurements are represented as the mean of 2 biological replicates +/-SD. **b**,**c**. Growth curves of PA14 WT and *ΔamrZ* infected with either a high (d,10^6^ PFU) or low (e, 10^4^ PFU) MOI of virulent DMS3m_*acrIF4*_. Phage replication was quantified as PFUs after 24 hours, and the efficiency of immunity expressed as a ratio of the number of PFUs harvested from *ΔamrZ* to the number of PFUs obtained from WT. OD600 and EOI measurements are represented as the mean of 3 biological replicates +/-SD. **d, e**. Timecourse of the fluorescence levels of Csy1-sfCherry reporter strains during surface-association (d) or planktonic growth (e). **f**. Overexpression of the indicated gene in a WT *csy1-sfCherry* background, with fluorescence measurements at 10 hours in liquid culture and normalized to empty vector. Fluorescence measurements are represented as the mean of 3 biological replicates +/-SD.

We next considered if phages and other mobile genetic parasites naturally antagonized by CRISPR-Cas immunity had evolved mechanisms to manipulate this repressive pathway. Inspired by the discovery of a *Paraburkholderia* phage that carried a distant homolog of AmrZ^59^, we searched the NCBI database for AmrZ homologs on *Pseudomonas* mobile genetic elements (MGE). Excitingly, we identified 14 diverse *Pseudomonas* genetic parasites carrying AmrZ homologs (Supplementary Table 2). These MGEs included obligately lytic and temperate Myoviridae, temperate Siphoviridae, and plasmids. AmrZ has been structurally characterized in complex with operator DNA^58^, and these mobile AmrZ homologs showed perfect conservation of critical DNA-interacting residues in the ribbon-helix-helix domain, suggesting conserved binding specificity (Fig. 4a, b, red residues/arrowheads). To test if these mobilized AmrZ variants were capable of regulating CRISPR-Cas activity in *Pseudomonas aeruginosa*, we synthesized 6 MGE-borne *amrZ* homologs and tested them for their ability to complement the *ΔamrZ* strain. 5/6 homologs complemented the *ΔamrZ* mutant, indicating they were active in the PA14 transcriptional network and were *bona fide* AmrZ orthologs (Fig. 4c). Next, each gene was expressed in WT cells carrying an endogenous *csy1-sfCherry* reporter, revealing 3 *P. aeruginosa* phage homologs (*amrZ*_*PaBG*,_ *amrZ*_*phi3*,_ *amrZ*_*JBD68*_) that inhibited Csy complex biogenesis at varying efficiencies (Fig. 4d). This suggests the possibility that *amrZ* has been hijacked for its capacity to inhibit CRISPR-Cas expression. Thus, *amrZ* acts on both sides of the battle between bacteria and mobile genetic elements.

**Figure 4.**
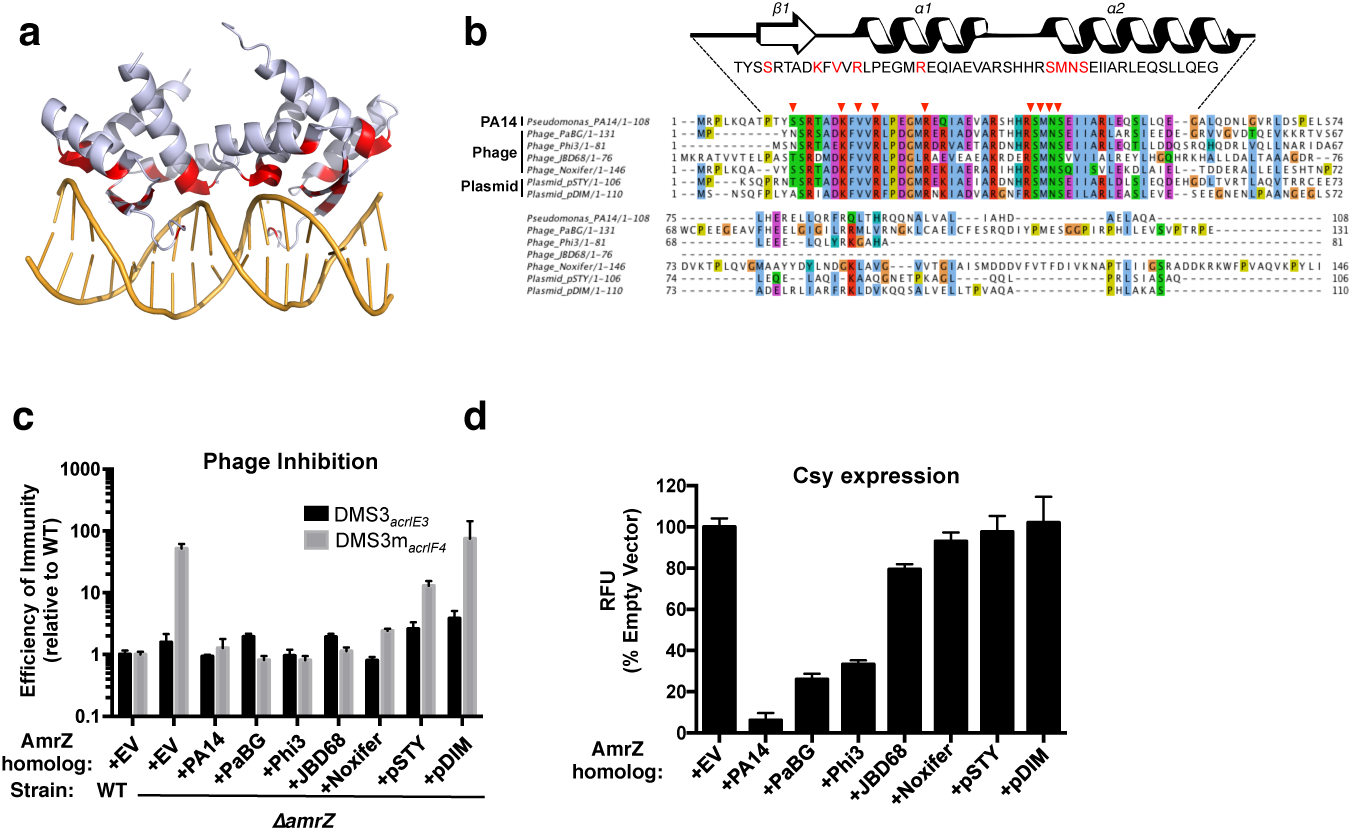
Phage AmrZ homologs control CRISPR-Cas immunity. **a.** Structure of an AmrZ tetramer bound to 18bp of operator DNA^58^ with DNA-contacting residues highlighted in red. **b.** Protein alignment of six mobile AmrZ homologs and the native PA14 AmrZ homolog, with the ribbon-helix-helix DNA binding domain secondary structure schematized and DNA-contacting residues indicated with red arrows and text. **c.** Efficiency of immunity (EOI) against isogenic phages DMS3_*acrIE3*_ (non-targeted) and DMS3m_*acrIF4*_ (CRISPR-targeted). Plaque forming units (PFUs) were quantified on ΔamrZ or the strains complemented with bacterial or MGE AmrZ homologs, and represented as a ratio to the number of PFUs measured on WT PA14. EOI measurements as represented as the mean of 3 biological replicates +/-SD. **d.** Measurement of the normalized fluorescence levels of Csy1-sfCherry reporter strains transformed with bacterial or MGE AmrZ homologs after 10 hours of growth in liquid culture. Fluorescence measurements are represented as the mean of 3 biological replicates +/-SD.

## Discussion

The factors that govern expression of bacterial processes are highly variable across species, reflecting niche-specific adaptations. CRISPR-Cas immune systems limit the replication of foreign genetic elements, but few reports address how the regulation of CRISPR-Cas impacts phage targeting. Here, we utilized a genetic screen that identified KinB as a positive regulator of CRISPR-Cas function in *P. aeruginosa.* This sensor kinase/phosphatase operates in a two-component system with AlgB that regulates a well-characterized biofilm-stimulatory pathway, though the ligand for KinB remains unknown. Removal of KinB or inactivation of its phosphatase activity leads to the accumulation of P-AlgB, which stimulates the activity of CRISPR-Cas repressors AlgU, AlgR, and AmrZ. This pathway is well-studied due to the recurrent isolation of alginate-overproducing (mucoid) *P. aeruginosa* from the lungs of cystic fibrosis patients, with mutations in *mucA*^60^ and *kinB*^47,61^ that lead to the constitutive activation of AlgU. This biofilm pathway generates the characteristic mucoidy phenotype of CF *P. aeruginosa* isolates^43,62,63^, which we also observed in *kinB:Tn* mutants (Supplementary Fig. 1a).

This pathway is well described in its capacity to activate biofilm production, and here we show that this transcriptional network also functions to dampen CRISPR-Cas activity in non-biofilm conditions. In WT cells, KinB phosphatase activity increases CRISPR-Cas function, though the effect is relatively mild in our experimental conditions. Relieving transcriptional repression via deletion of *algU* or *algR* however leads to a hyper-activation of the CRISPR-Cas immune response. We further demonstrate that *amrZ* acts in this pathway to specifically repress CRISPR-Cas when cells are surface-associated, leading to lower levels of Csy complex during surface-association relative to planktonic growth. Phages and plasmids have taken advantage of this regulatory scheme, and some *Pseudomonas* genetic parasites carry hijacked copies of the CRISPR-Cas repressor *amrZ*. We show that these mobilized *amrZ* homologs retain their capacity to transcriptionally repress CRISPR-Cas, demonstrating a dual role for this gene in the host-pathogen arms race between bacteria and their parasites.

Considering the physiology of *P. aeruginosa* can put these findings into context with a phage and CRISPR-Cas-centric view. During exponential growth is when we and others observe CRISPR-Cas activation^8,9^, and is also when phage infection risk is likely high (i.e. metabolically active, well-mixed planktonic culture^33^). Conversely, surface-association lessens phage infection risk, as spatial structure limits phage dispersal and prevents a phage bloom from overtaking the entire bacterial population^64^. Though we do not measure phage sensitivity in a biofilm here, biofilm production likely provides an even greater level of intrinsic phage resistance, due to high levels of spatial stratification and abundant polysaccharide secretion.

Supporting our proposal that *P. aeruginosa* surface-association/biofilm formation and CRISPR-Cas expression are inversely correlated are two independent observations: **i)** Our analysis of a previously published PA14 RNAseq data set^65^ and proteomic data set^66^ revealed activation of CRISPR-Cas expression in exponential phase, and strong transcriptional repression during stationary phase and biofilm growth (Supplementary Fig. 4a). Interestingly Cas proteins are still detected in stationary phase and biofilm growth, suggesting the cells retain some immune-competency even after transcriptional shutdown (Supplementary Fig. 4b). **ii)** Previous studies on the *P. aeruginosa* CRISPR-Cas system have revealed that the genome is hyper-sensitive to CRISPR-induced DNA damage during surface-association and biofilm formation, leading to cell death when a mismatched prophage sequence target is present in the chromosome^30,31^. This suggests that CRISPR auto-immunity costs are also dependent on the growth state and physical environment of the cell, and that CRISPR-Cas regulation in *P. aeruginosa* is tuned to reflect this intrinsic sensitivity.

Here, we report the identification of a CRISPR-Cas repressive pathway in *P. aeruginosa*. Phages bearing distinct anti-CRISPRs are employed in bioassays to read out the activity of the CRISPR-Cas surveillance complex. Using a CRISPRi phage-repression assay, coupled with translational reporters, we conclude that *algU, algR*, and *amrZ* control the production of the CRISPR surveillance complex (Csy complex), which is where the screen was focused. Cas3 levels, though low, are also modulated by the KinB/AlgB pathway. Interestingly, our observations show that CRISPR-Cas activity in *P. aeruginosa* does not operate at full strength, despite the capacity of a hyper-activated immune system to severely limit the replication of phages relying on anti-CRISPRs for survival. We speculate that the ability to control CRISPR-Cas activity during lifestyle transitions may be essential for *P. aeruginosa* to safely maintain a CRISPR-Cas system by limiting self-toxicity. Furthermore, we reveal an unexpected cost to negative regulation of CRISPR-Cas immunity in our discovery of MGE-encoded CRISPR-Cas repressors: though CRISPR-Cas control is likely essential for safe retention of the immune system, the evolution of potent mechanisms of CRISPR-Cas repression has created an Achilles Heel that is exploited by genetic parasites.

## Acknowledgements

We thank members of the Bondy-Denomy lab for conversations and support relating to this project. We also thank Kristine Trotta (Seemay Chou’s lab, UCSF) for assistance and advice in the development of fluorescent reporter assays, and Andrew Santiago-Frangos (Blake Wiedenheft’s lab, MSU) for advice and consultation. The Bondy-Denomy lab was supported by the University of California, San Francisco Program for Breakthrough in Biomedical Research, funded in part by the Sandler Foundation, and an NIH Office of the Director Early Independence Award (DP5-OD021344), and R01GM127489. We thank Deborah Hung’s lab for providing *ΔkinB, ΔalgR, ΔalgU* strains and associated mutants, and George O’Toole’s lab providing the *csy3::lacZ* PA14 strain.

## Author Contributions

J.B.D., A.L.B, and B.C. formulated study design and plans. A.L.B. designed and conducted bacteriophage plaque assays, liquid infection assays, and CRISPRi assays, performed CRISPR-Cas expression profiling, and conducted bioinformatics analyses. B.C. conducted the genetic screen, isolated and constructed bacterial mutants, conducted LacZ expression profiling, and performed bacteriophage plaque assays. S.G. designed, constructed, and validated sfCherry reporter strains. T.S. conducted CRISPRi assays and V.E. assisted in establishing fluorescent reporter assays. J.B.D. and A.L.B wrote the manuscript.

## Materials & Correspondence

Correspondence should be directed to

## Methods

### Bacterial strains and bacteriophages

*P. aeruginosa* UCBPP-PA14 (PA14) strains and *E. Coli* strains (Sup. Table) were grown on lysogeny broth (LB) agar or liquid at 37°C. Media was supplemented with gentamicin (50 µg ml^-1^ for *P. aeruginosa* and 30 µg ml^-1^ for *E. Coli*) to maintain the pHERD30T plasmid or carbenicillin (250 µg ml^-1^) for *P. aeruginosa* or ampicillin (100 µg ml^-1^) for *E. coli* containing the pHERD20T plasmids. pHERD plasmids were induced with 0.1% arabinose. Bacteriophages stocks were prepared as described previously^40^. In brief, 3 ml of SM buffer was added to plate lysates of the desired purified bacteriophage and incubated at room temperature for 15 minutes. SM buffer containing phages was collected and 100 µl of chloroform was added. This was centrifuged at 10,000 x g for 5 minutes and supernatant containing phages was transferred to a screw cap storage tube and incubated at 4 °C.

### Tn mutagenesis screen

The *csy*::*lacZ* reporter strain was subjected to transposon mutagenesis and colonies were isolated on plates containing X-gal (5-bromo-4-chloro-3-indolyl-β-D-galactopyranoside). ∼50,000 colonies were visually examined for increased or decreased levels of β-galactosidase and insertions mapped by semi-random PCR. To conduct transposon mutagenesis, overnights of PA14 *csy3::lacZ* and *E. coli* containing the pBTK30 Tn suicide vector were mixed in a 1:2 ratio (donor : recipient) for conjugation. A second control group containing PA14wt and *E. Coli* containing the pBTK30 Tn suicide vector was set up in parallel. Mixed cells were centrifuged at 4,000 x g for 10 minutes to pellet cells. 100 µl of resuspended conjugation pellet was then spotted on LB agar plates and incubated at 37 °C for 6hrs. Conjugation spots were collected and resuspended in LB liquid media. Conjugation was then screened on an LB agar plates supplemented with nalidixic acid (30 µg ml^-^1) and gentamicin (50 µg ml^-1^). Surviving colonies containing Tn insertions were collected into 1ml of LB liquid media. Serial dilution of were prepared and plated on LB agar plates supplemented with x-gal (200 µg ml^-1^) and gentamicin (50 µg ml^-1^) and nalidixic acid (30 µg ml^-^1). Plates were incubated at 37 °C for 24 hours to allow for colonies to change color. Colonies displaying changed expression levels as compared to our positive control (PA14 csy3::lacZ no pBTK30) were then isolated onto secondary LB agar plates with X-gal and gentamicin and nalidixic acid at the stated concentrations. Genomic DNA (gDNA) was collected from isolated single colonies by resuspending bacterial colonies in 0.02% SDS and boiling the sample for 15 minutes. Samples were then centrifuged at 10,000 x g and supernatants containing gDNA were collected. Semi-random PCR was then performed using PCR primers listed on sup table_. PCR samples were sequenced and reads were then matched to the *P. aeruginosa* UCBPP-PA14 genome using BLAST. Isolated containing genes of interest were saved as stocks (s. table 1). Transcription changes were then verified via modified β-galactosidase assay.

### Plaque assays

Plaque assays were performed on LB agar plates (1.5% agar) with LB top agar (0.7% agar), both supplemented MgSO_4_ (10 mM final concentration) and gentamicin (50 µg ml^-1^) as needed for plasmid maintenance. Spot titrations were done by mixing 150 µl of a *P. aeruginosa* overnight culture with 3 ml of top agar, which was dispersed evenly on a LB MgSO_4_ plate. 3 µl of 10-fold phage dilutions were then spotted on the surface. Plates were incubated overnight at 30 °C. To count plaques, full plate assays were used, except when CRISPR-targeting was so strong that discrete plaques could not be accurately measured. In this case, spot titrations are shown. For full plate assays, 3 µl of the selected phage dilution was incubated with 150 µl of *P. aeruginosa* overnight for 15 minutes at 37 °C. 3 ml of top agar was then added and the mixture was dispersed evenly on a LB MgSO_4_ plate. Individual plaques were then counted to assess differences in plaquing efficiency.

### β-galactosidase assay

A modified version of the previously described β-galactosidase assay was used to measure *lacZ* activity in transcriptional fusions. Bacterial cultures were grown overnight at 37 °C. Cultures were then diluted 1:100 into LB liquid medium supplemented with the desired antibiotic. Diluted cultures were then incubated at 37 °C until the desired time point was reached. Culture density was measured with a spectrophotometer (OD_600_) and 200 µl of the sample was added 800 µl to permeabilization solution (components listed in sup table XX). Cells were mixed via inversion and vortexed for 1 minute to permeabilize the cells. 200 µl of ONPG (4 mg ml^-1^) was added and samples were incubated for 27 minutes at 30 °C. Enzymatic reaction was stopped by addition of 300 µl of 1M Na_2_CO_3_. Samples were centrifuged at 13,000 x g for 5 minutes to remove debris and 200 µl of supernatant was moved to a 96-well plate to read absorbance at 420 nm and at 550 nm. Miller units were calculated using the Miller equation. 3 technical replicates per sample per experiment.

### Phage transduction of *kinB::Tn* alleles

Transposon insertions in *kinB* from a *csy3::lacZ* background were transduced into WT PA14 to enable testing of CRISPR-Cas function with the same transposon insertion. Phage phiPA3 was used to infect the donor strain (*csy3::lacZ, kinB::Tn*), on plates with top agar overlays, using ∼10^4^ PFU to generate near confluent lysis. Plates were soaked in 3-4 mL of phage SM buffer and 2 mL collected over chloroform, vortexed, and pelleted to isolate transducing phage in the supernatant. Lysates were used to infect recipient strains (WT PA14). ∼10^8^ PFU were used to infect a culture at an MOI of 1. After 30 minutes of static incubation on the bench, cultures were gently shaken at 37 °C for 20 min and then pelleted at 5000 x g. Cells were washed twice with LB, and subsequently incubated at 37 °C for 1 hour to allow recombination and gentamicin resistance outgrowth. Cultures were pelleted and resuspended in 200 µL of LB, and plated on LB plates containing gentamicin. Controls included uninfected cells and cells infected with phages not propagated on a gentamicin resistant donor strain. Additionally, phage lysate was directly plated under selection to confirm no residual donor strain in the phage preparation. Plates were incubated overnight at 37 °C and their identity (i.e. CRISPR-Cas intact) confirmed with a plaque assay using DMS3m as the target phage and PCR of the kinB locus.

### Introduction of *csy3::lacZ P. aeruginosa* UCBPP-PA14 strains

The *lacZ* gene was introduced into PA14 strains of interest via homologous recombination. Recombination vector containing lacZ flanked by homology arms matching csy2 and csy4 was introduced via conjugation. PA14 strains and *E. coli* containing vector were mixed at a ratio of 1:2 (recipient:donor). Mixture was heat shocked at 42 °C for 10 min. Mating spot was then plated on a LB agar plate and incubated overnight for 30 °C. Mating spot was then collected, resuspended in 1 ml of LB liquid media and plated on VBMM plates supplemented with 50 ug/mL gentamicin to select for colonies with the integrated homology plasmid. Colonies were cultured overnight in LB in the absence of selection at 37 °C, and were then diluted and counterselected on no salt LB (NSLB) agar plates supplemented with 15% sucrose. Surviving colonies were then grown on LB agar plates supplemented with gentamicin and X-gal to check for *lacZ* insertion via color change and lacZ insertion was further verified via PCR.

### Generation of an PA14*δamrZ* strain using the endogenous I-F CRISPR-Cas system

Complementary oligonucleotides encoding a crRNA targeting the *amrZ* gene of PA14 were annealed and ligated into the multiple cloning site of the pHERD30T vector. A fragment possessing homology arms flanking the desired mutation (500 bp upstream and 500 bp downstream) around *amrZ* was cloned into a distinct location (NheI site) of the same vector via Gibson assembly. The new plasmid containing both a crRNA and homology region was introduced into WT PA14 via electroporation. Transformation efficiency dropped dramatically in the presence of the crRNA due to the toxicity caused by self-targeting. All surviving colonies had the desired clean deletion of the *amrZ* gene. Deletions were confirmed by PCR of the region of interest and subsequent Sanger sequencing of the amplicon. A 2000 bp region flanking *amrZ* was PCR amplified and sequencing primers were designed to sequence both the deletion junction and outside of the original 500 bp flanking regions to confirm the removal of the *amrZ* gene.

### Liquid phage cooperation assay

Liquid phage infections were performed as described in^41^. In brief, an overnight culture of cells was diluted 1:100 into fresh media, and infected with virulent phage DMS3m_*acrIF4*_ in biological triplicate in a 96 well Costar plate. Cells were incubated at 37 °C with constant rotation and OD600 measured every 5 minutes in a Biotek H4 Synergy plate reader. Phage were harvested from each well and quantified by plaque assay after 24 hours. In these experiments, all strains used in the assay carried 2 spacers against the DMS3m_*acrIF4*_ phage to prevent phage escape: one endogenous spacer (CRISPR2_sp1), and the other spacer was provided on a pHERD30T plasmid.

### Generation of endogenous Csy1-sfCherry and Cas3-sfCherry reporters

Endogenous Csy1-sfCherry and Cas3-sfCherry reporters were constructed similar to the construction of *csy3::lacZ.* We initially verified that tagging of sfCherry at the N terminus of Csy1 and Cas3 are functional, when expressed from a plasmid. pMQ30-sfCherry-Csy1, which contains sfCherry sequence flanked by 657 bp upstream of csy1 and 701 bp downstream of *csy1* start codon, was cloned in pMQ30 plasmid between HindIII and BamHI sites using Gibson assembly. pMQ30-sfCherry-Cas3, which contains sfCherry sequence flanked by 353 bp upstream of cas3 and 350 bp downstream of cas3 start codon, was cloned in pMQ30 plasmid between HindIII and BamHI sites using Gibson assembly. 4 bp that overlap between the end of *cas1* and the beginning of *cas3* were duplicated in the final construct. Both pMQ30-sfCherry-Csy1 and pMQ30-sfCherry-Cas3 contains ggaggcggtggagcc sequence (encoding GGGGA) as linker between sfCherry and the respective tagged proteins. The Csy1-sfCherry and Cas3-sfCherry construct were introduced into PA14 strains of interest via homologous recombination. Strains containing appropriate insertion were verified via PCR.

### sfCherry reporter profiling. Liquid

Cells were diluted 1:100 from an overnight culture into fresh LB (with antibiotics and inducer if required), and grown for the indicated number of hours in biological triplicate. 500 µl of each sample was then spun down at 8,000xg for 2 minutes, and resuspended in 500 µl of M9 media. Samples were loaded in to a 96 well plate (150 µl /well) in technical triplicate and red fluorescence and OD600 were measured using a Biotek H4 Synergy. **Solid:** Cells were diluted 1:100 from an overnight culture into fresh LB and 20 µl plated onto individual wells in biological triplicate in a 24 well plate with each well containing solidified 1.5% LB Agar with antibiotics and inducer if required. The 24 well plate was then covered with a breathable Aeraseal, and incubated at 37 °C with no shaking. At the indicated timepoint, cells were harvested by flooding each well with 500 µl of M9 buffer, and were spun down at 8,000xg for 2 minutes, and resuspended in 500 µl of M9 media. Samples were loaded in to a 96 well plate (150 µl /well) in technical triplicate and red fluorescence (excitation 580 nm, emission 610 nm) and OD600 were measured using a Biotek H4 Synergy. To calculate the relative fluorescence units for each sample, the background fluorescence and background OD600 values obtained were subtracted from the sample values, and the sample fluorescence was then normalized to the sample OD600.

### AmrZ homolog discovery and characterization

BLASTp was used to search the nonredundant protein database for AmrZ homologs (accession: ABJ12639.1) in *Pseudmonas sp.* (taxid: 286) in May 2019. This homolog list (e value > 0.001) was then examined for homologs found on phage or plasmid genomes. Representative homologs were aligned using Clustal and the alignment visualized in Jalview, and key conserved residues were mapped onto the structure in Pymol (PDB ID: 3QOQ). Select homologs were synthesized (TWIST Biosciences) and cloned into the SacI/PstI site of the arabinose-inducible plasmid pHERD30T using Gibson assembly. Vectors were electroporated into *Pseudomonas aeruginosa* strains for functional testing, where they were induced with 0.1% arabinose and maintained with 50 ug/mL gentimicin.

---

**Supplementary Table 1.** All independent transposon insertions identified and mapped by visual screening with increased or decreased *csy3::lacZ* β-galactosidase activity at 8 hours of growth. The insertion location in the PA14 genome is shown, along with the measured level of β-galactosidase enzyme. These measurements were not determined (N/D) for strains with a growth defect.

| Gene with Tn insertion | Transposon location | β-gal activity (% of unmutagenized) |
| --- | --- | --- |
| kinB (1) | 6448519 | 59% |
| kinB (2) | 6447945 | 50% |
| kinB (3) | 6447811 | 53% |
| kinB (4) | 6449373 | 54% |
| kinB (5) | 6449345 | 98% |
| purM | 4618060 | 134% |
| lasR | 4085810 | 135% |
| minD | 1915545 | N/D (growth defect) |
| gltB | 5943637 | 108% |
| S-type pyocin | 1196879 | 145% |
| pyoS3A | 4404303 | 145% |
| tolA | 4595505 | 158% |
| glycosyl transferase | 5889967 | 81% |
| bacA | 3490006 | 75% |
| Intergenic; zipA and smc + | 3979746 | N/D (growth defect) |
| gidB | 6530337 | 88% |
| deaD | 2373899 | N/D (growth defect) |
| polA | 6457284 | N/D (growth defect) |
| pchH | 798159 | 131% |
| crc | 6275250 | 91% |
| putative plasmid stabilization protein | 5347104 | N/D (growth defect) |
| paraquat inducible protein | 5532785 | 108% |
| pchH | 797705 | 127% |
| yhiH/yhil | 6162134 | 108% |
| oxidoreductase FMN binding | 4188602 | N/D (growth defect) |
| putative membrane protein | 1400718 | 118% |
| Intergenic; fstA and fstZ | 5104077 | 111% (growth defect) |
| putative Zn-dependent oxidoreductase | 2820446 | 115% (growth defect) |
| gnyL | 3434168 | 87% |
| cytochrome c1 precursor | 5126449 | 94% |

**Supplementary Table 2:**
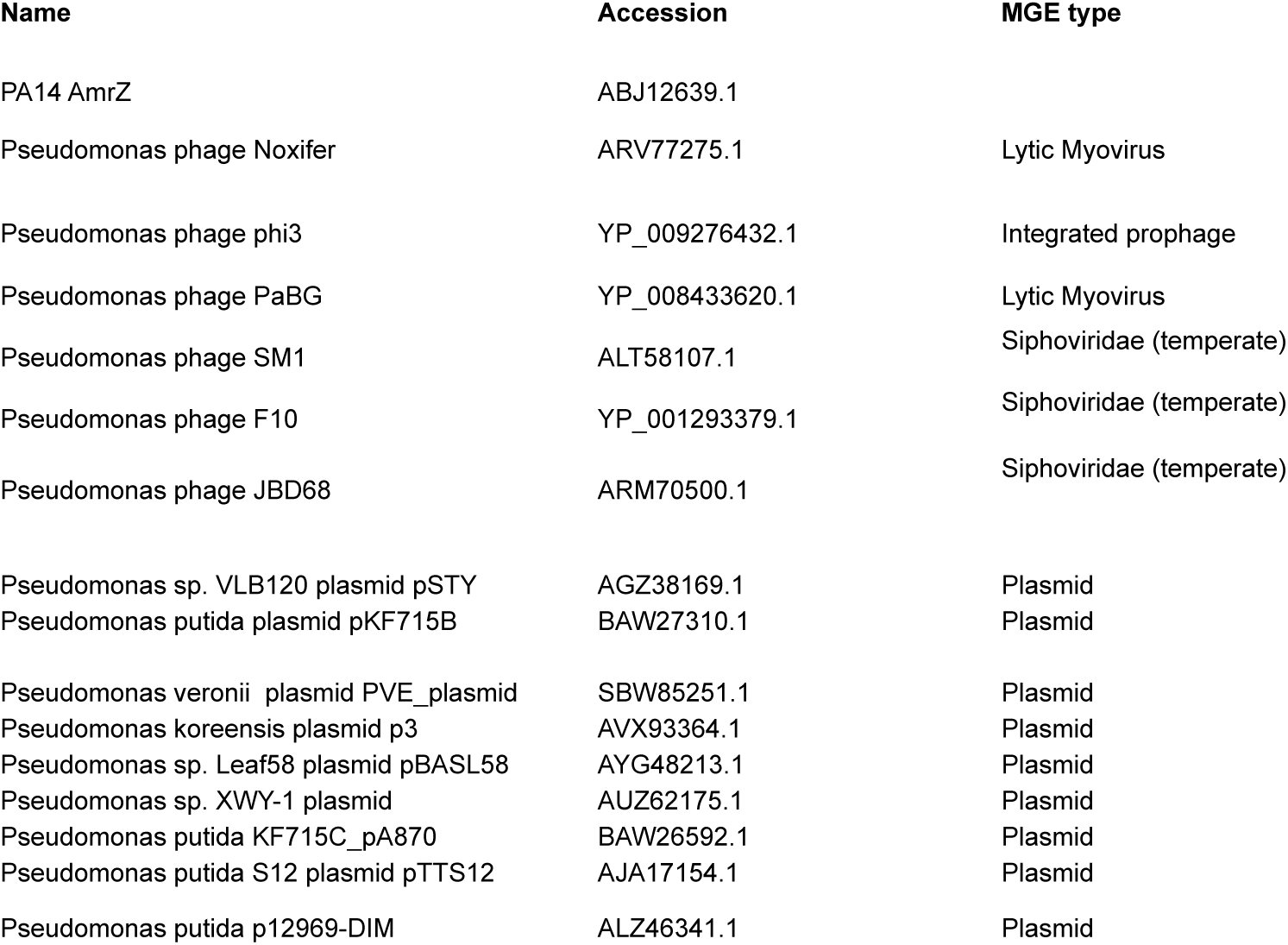
AmrZ homologs listed by the genome that encodes them, the accession number, and the mobile genetic element type.

**Supplementary Figure 1.**
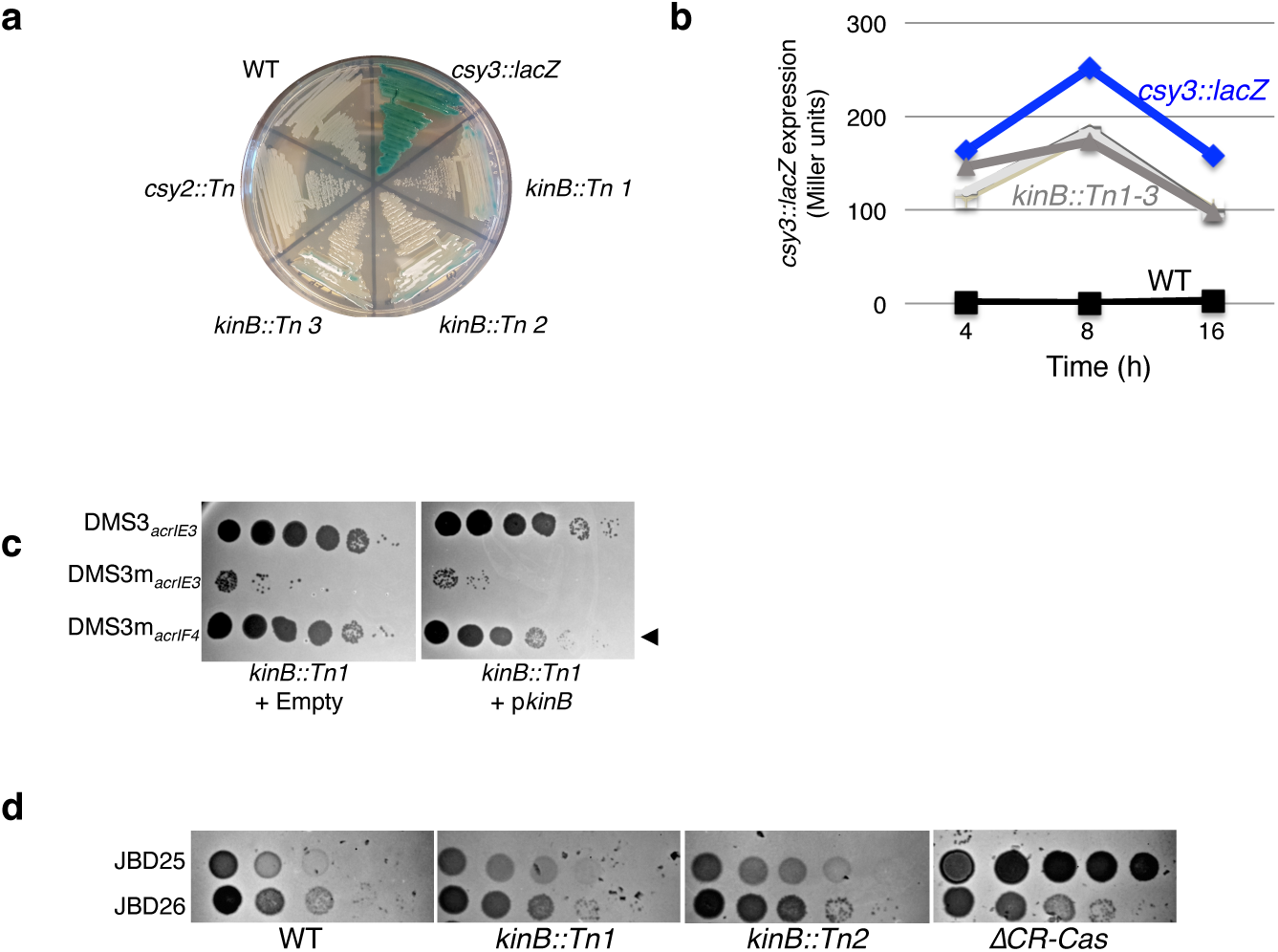
**a.** A streak plate on X-gal plates, showing strains involved in this study and isolated transposon (Tn) insertions. **b**. β-galactosidase measurements at the indicated time points for the unmutagenized (*csy3::lacZ*) strain and three isolated *kinB* transposon mutants (*kinB::Tn1-3*). **c**. Phage titration on lawns of the *kinB::Tn1* mutant transformed with empty vector or *kinB.* **d.** Spot titration of phages JBD26 (CR2_sp17, sp20-targeted, possessing *acrIF4*), JBD25 (CR1_sp1 targeted) on *kinB::Tn* mutants and ΔCRISPR-Cas.

**Supplementary Figure 2:**
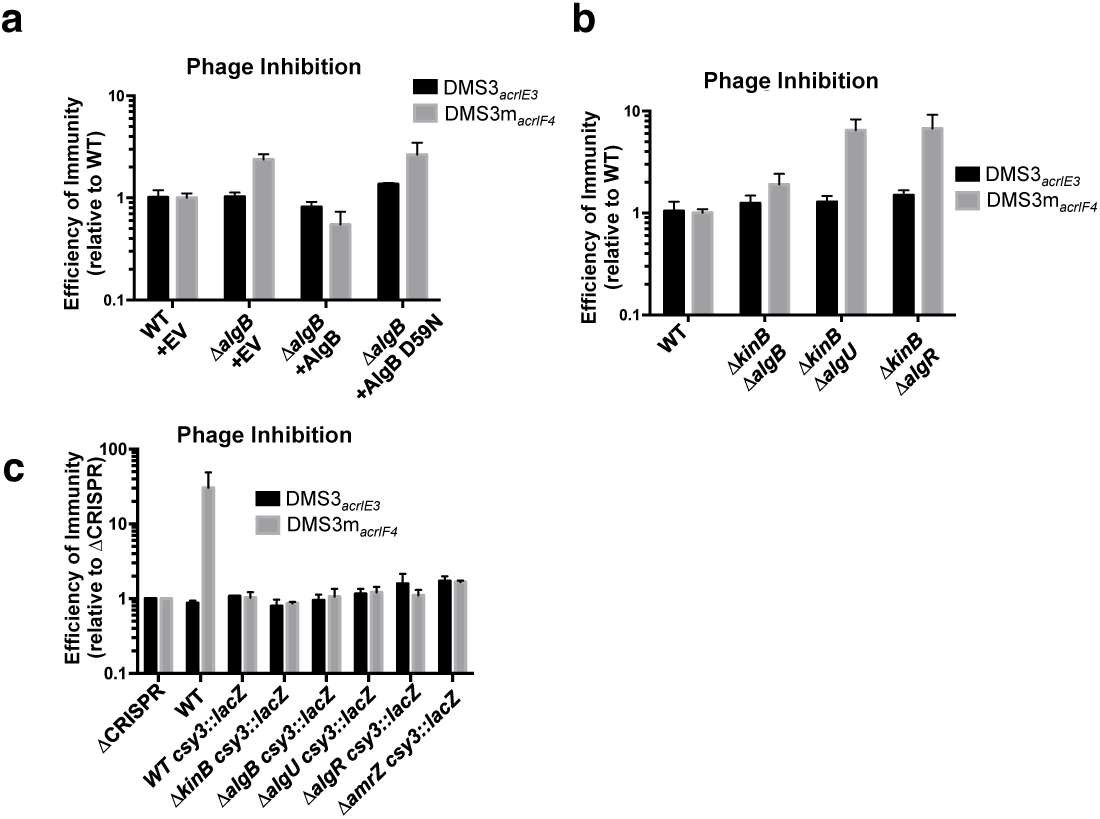
**a-c.** Efficiency of immunity measurements for indicated mutants relative to WT. **a**. *ΔalgB* mutant complemented. **b.** Double knockouts show *ΔkinB* combined with *algB, algU*, or *algR*. **c**. Indicated knockouts were combined with *csy3::lacZ*.

**Supplementary Figure 3:**
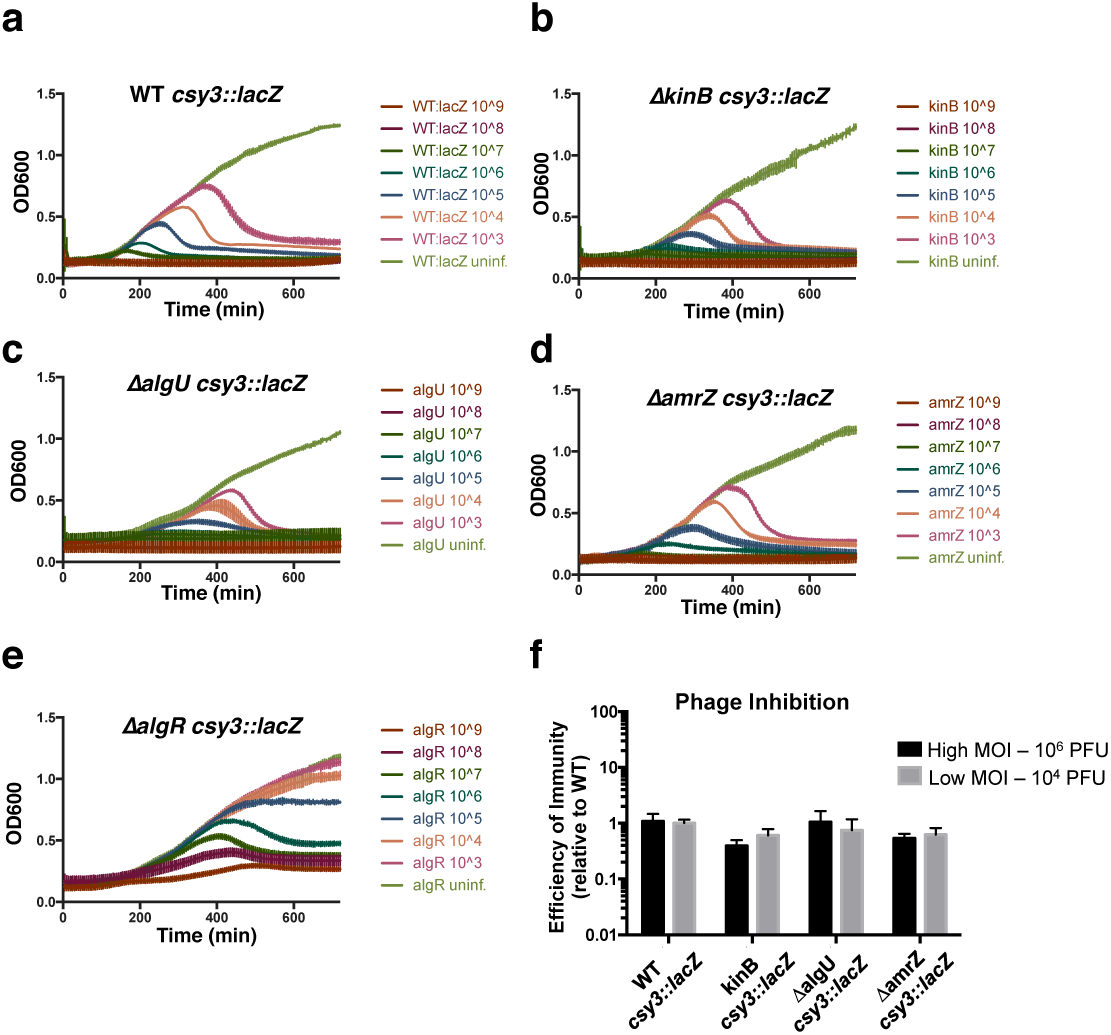
**a-e**. Growth curves of the indicated knockouts combined with *csy3::lacZ*. infected with virulent DMS3m_*acrIF4*_ at multiplicity of infection (MOI, rainbow colors) increasing in 10-fold steps from 2×10^-4^ to 2×10^2^. **f.** Phage replication was quantified as PFUs after 24 hours for the MOIs used in Figure 1 and 3, and the efficiency of immunity expressed as a ratio of the number of PFUs harvested from the mutants combined with *csy3::lacZ* under the number of PFUs obtained from WT *csy::lacZ*. OD600 and EOI measurements are represented as the mean of 3 biological replicates +/-SD.

**Supplementary Figure 4.**
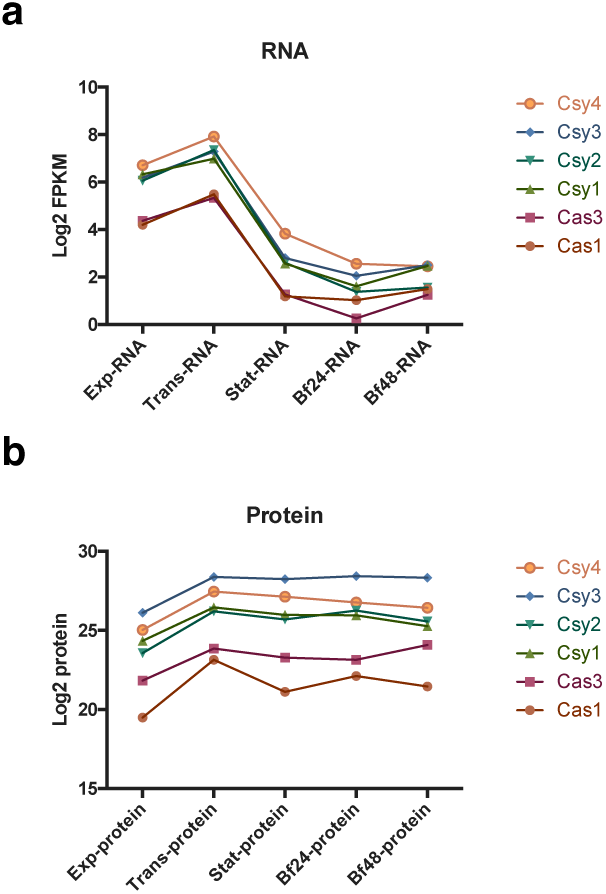
**a.** Log2 of Fragments Per Kilobase of transcript per Million mapped reads (FPKM) shown for each I-F *cas* gene in PA14 in the indicated growth condition. **b.** Log2 of protein levels for each of the I-F Cas proteins in PA14 in the indicated growth condition.

